# Machine learning based classification of cells into chronological stages using single-cell transcriptomics

**DOI:** 10.1101/303214

**Authors:** Sumeet Pal Singh, Sharan Janjuha, Samata Chaudhuri, Susanne Reinhardt, Sevina Dietz, Anne Eugster, Halil Bilgin, Selçuk Korkmaz, John E. Reid, Gökmen Zararsiz, Nikolay Ninov

**Author notes:** Correspondence to: sumeet and.

## Abstract

Age-associated deterioration of cellular physiology leads to pathological conditions. The ability to detect premature aging could provide a window for preventive therapies against age-related diseases. However, the techniques for determining cellular age are limited, as they rely on a limited set of histological markers and lack predictive power. Here, we implement GERAS (GEnetic Reference for Age of Single-cell), a machine learning based framework capable of assigning individual cells to chronological stages based on their trans criptomes. GERAS displays greater than 90% accuracy in classifying the chronological stage of zebrafish and human pancreatic cells. The framework demonstrates robustness against biological and technical noise, as evaluated by its performance on independent samplings of single-cells. Additionally, GERAS determines the impact of differences in calorie intake and BMI on the aging of zebrafish and human pancreatic cells, respectively. We further harness the predictive power of GERAS to identify genome-wide molecular factors that correlate with aging. We show that one of these factors, *junb*, is necessary to maintain the proliferative state of juvenile beta-cells. Our results showcase the applicability of a machine learning framework to classify the chronological stage of heterogeneous cell populations, while enabling to detect pro-aging factors and candidate genes associated with aging.

## BACKGROUND

Aging is a universal phenomenon, during which cells undergo progressive transcriptional ^1,2^, genomic ^3,4^ epigenetic ^5^, and metabolic ^6^ changes. The age-related modifications can deteriorate the functional properties of cells. The accumulation of cellular defects can lead to a decline in organismal health and to the onset of age-related diseases. A major focus of the biology of aging is to identify factors that accelerate or slow-down, preferably even reverse, the cellular aging process. Biological studies have identified multiple modifiers of the aging process, including genetic and environmental factors ^7,8^. For instance, caloric restriction has been demonstrated to increase lifespan in multiple species ^9^, including humans ^10^. However, the discovery of factors that influence aging relies on retrospective measures, after the impact of age has already manifested itself, and depends on a restricted set of indicators based on histological analysis ^11^. It is therefore imperative to develop reliable indicators of cellular age that forgo the need for detrimental phenotypes. Predicting cellular aging before the defects manifest themselves would provide a window for therapeutic interventions. Preventive therapies during this window would bypass additional complications arising after the onset of the pathology.

The development of a reliable cellular age predictor requires two principal components. Firstly, it entails a reliable assessment of the transitions cells undergo with age. Secondly, the predictor should be capable of placing cells of unknown age along this transition path in order to estimate their age. The first objective, assessment of cellular transitions, has been enabled by recent advances in single-cell mRNA expression profiling ^12^. Cellular progression through the transitions is increasingly being described by both heuristic methods and probabilistic models. These methods are categorized as pseudotemporal estimation algorithms and use techniques such as dimensionality reduction, graph theory, bifurcation analysis and optimal-transport analysis to place cells along a transition trajectory ^13-18^. All the methods make explicit or implicit assumptions about the smoothness of mRNA expression profiles along the trajectories and seek to explain part of the variation across the cells by location along the trajectory. Unwanted variation that cannot be explained by trajectory location can confound the analysis. Some methods protect against confounding effects by using a prior over pseudotime that leverages information about the time cells were assayed ^18^ whilst others do not. Although current methods can reveal cellular transitions during a differentiation process ^19-22^, they have only been shown to work retrospectively, that is they have no predictive ability to insert *de-novo* samples into the trajectories. Thus, their predictive utility on unseen cells, the second objective, remains unresolved.

Prediction of the position of *de-novo* samples in a cellular transition trajectory requires discrimination of the transcriptional features of importance from the confounding factors that accompany single-cell measurements. The three main confounding factors are: 1) biological noise due to fluctuations in mRNA expression levels, 2) technical noise inherent in single-cell mRNA sequencing, and 3) cell-type diversity within an organ. Biological noise can arise due to the stochasticity in biochemical processes involved in mRNA production and degradation^23,24^, heterogeneity in the cellular microenvironment^25^, and many more unknown factors. Although mechanisms such as the passive transport of newly transcribed mRNA from the nucleus to the cytoplasm exist to reduce the level of biological noise ^26^, it can never be eliminated completely ^23^. In fact, aging might enhance fluctuations in mRNA expression levels ^27,28^. Nevertheless, in certain contexts, fluctuations in expression levels are beneficial to the organism ^29,30^. Technical noise, on the other hand, arises due to the sensitivity and depth of single-cell sequencing technology ^31^. Sequencing involves conversion of mRNA into cDNA and amplification of the minute amounts of cDNA. These steps could omit certain mRNA molecules, muting their detection. Moreover, amplified cDNA molecules might escape sequencing due to the limits on the comprehensiveness of the technology. In effect, expression noise is inherent to single-cell measurements.

The diversity in cell types within an organ adds a second layer of complexity to the inherent noise in mRNA expression. Diverse types of cells express unique sets of genes and regulatory networks. Moreover, numerous studies have demonstrated the presence of cellular sub-populations within nominally homogenous cells ^32,33^. For example, pancreatic beta-cells have been shown to consist of dynamic sub-populations with different proliferative and functional properties ^34-36^, and liver cells were demonstrated to display variability in gene expression depending on their location within the organ ^37^ Thus, the inherent cell-to-cell heterogeneity adds to the challenge of extracting age-specific transitions from mRNA expression profiles. Furthermore, cellular heterogeneity makes it difficult to extrapolate the results from studies at the tissue-scale to the aging of individual cells and to identify common molecular signatures of aging ^38,39^.

In this study, we provide a framework that efficiently ‘learns’ the cellular transitions of aging from single-cell gene expression data in the presence of expression noise and cellular heterogeneity. First, the age predictor is trained to recognize the age of individual cells based on their chronological stage. Chronological stage is an easily measurable fact, and hence provides a ground truth for the training. Second, we show that the trained predictor can place robustly cells of unknown ages along the aging path. To show the utility of the age predictor, we apply it to the pancreatic beta-cells, which represent an excellent system for studying aging. In mammals, the beta-cell mass is established during infancy and serves the individual throughout life ^40^. The long-lived beta-cells support blood glucose regulation, with their dysfunction implicated in the development of Type 2 diabetes. Older beta-cells display hallmarks of aging, such as a reduced proliferative capacity and impaired function ^41^. We first focus on the zebrafish beta-cells due to the potential for visualization and genetic manipulation of beta-cells at single-cell resolution ^36^, and extend our framework to human pancreatic cells using publicly available published datasets. Finally, we demonstrate the predictor’s utility in identifying age-modifying genetic and environmental factors.

## RESULTS

### Machine learning based framework accurately and robustly predicts chronological stage

To capture the transcriptional dynamics of beta-cells with age, we performed singlecell mRNA sequencing of beta-cells in primary islets dissected from animals belonging to three chronological stages: Juvenile (1 month post-fertilization (mpf)), Adolescent (3, 4 and 6 mpf) and Adult (10, 12 and 14 mpf). Using *Tg(ins:Betabow)* ^36^, a transgenic line that specifically marks zebrafish beta-cells with red fluorescence (Supplementary Fig. S1), we isolated and sequenced 827 beta-cells in multiple batches. Sequencing was performed using the Smart-Seq2 protocol, which has been demonstrated to provide higher transcriptional coverage than other methods ^42^. The sequenced cells were quality-controlled to yield a total of 645 beta-cells (Supplementary Fig. S2). To identify age-specific transitions, we first attempted to order the cells using an unsupervised pseudotemporal analysis (Supplementary Fig. S3). However, the beta-cells from the three chronological stages were broadly spread along the predicted temporal trajectory. The shortfall of unsupervised pseudotemporal ordering prompted us to consider an alternative approach in which we modeled the data using the ground truth provided by the chronological stage. For this, we developed a supervised deep learning framework to predict the stage of the cellular origin: Juvenile, Adolescent or Adult (Fig. 1a). As input to the classifier, genes detected in all the cells were ranked in descending order of their variability and the top 1000 genes were selected for training (Supplementary Table S1). Since neural networks are prone to overfitting, two normalizing hyperparameters were added: L2 regularization (which penalizes a strong focus on few inputs) and dropout regularization (which helps ‘averaging’ across connections). This framework was named GERAS (GEnetic Reference for Age of Single-cell) in reference to the Greek God of old age.

**Figure 1:**
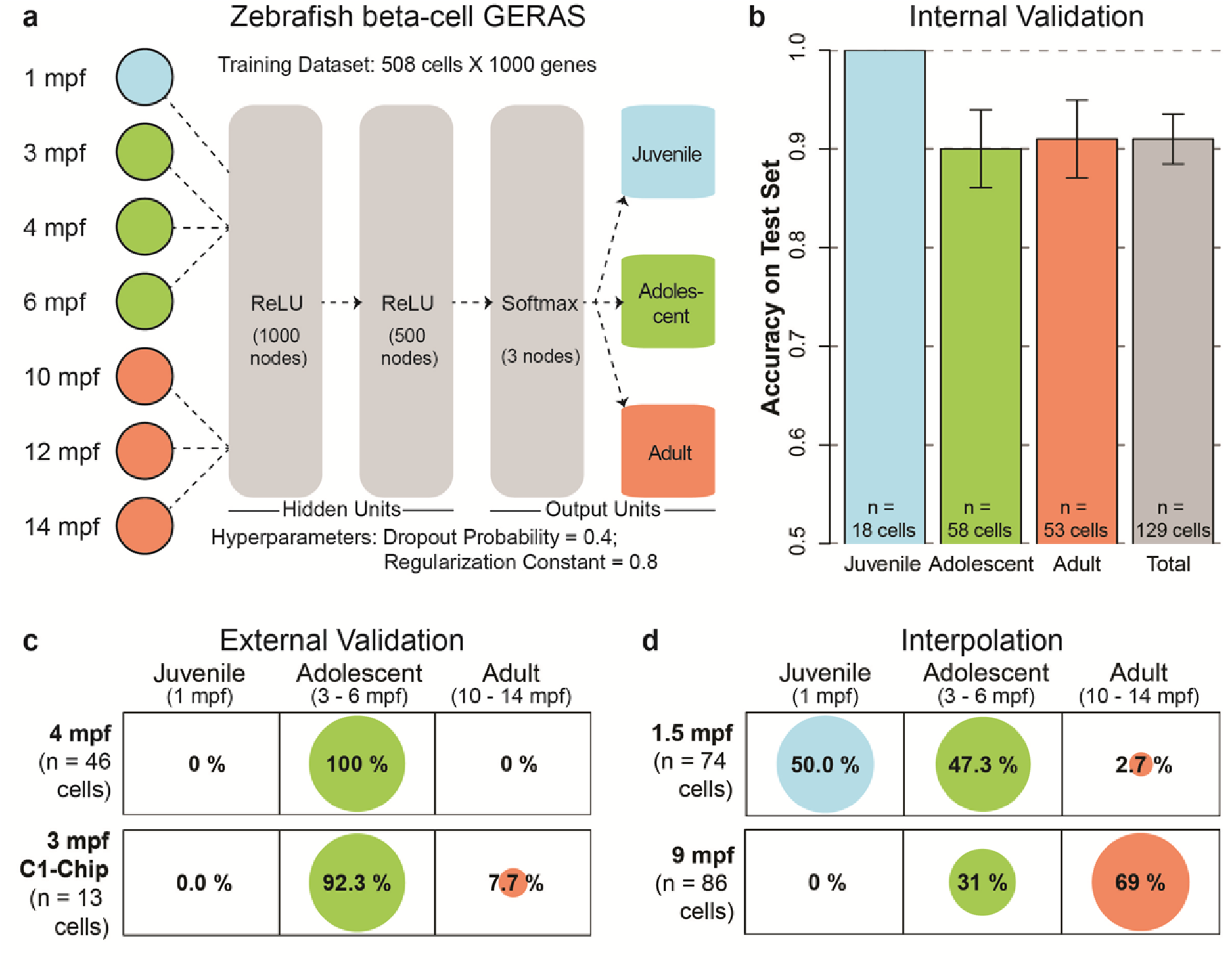
A Chronological age classifier for zebrafish beta-cells. **(a)** A schematic of the machine learning framework for predicting the chronological age of zebrafish beta-cells based on single-cell transcriptome (see Online Methods for details). **(b)** Barplot showing the accuracy of GERAS for classifying the age of beta-cells that were excluded during the training of the model. The predictions on the excluded beta-cells displayed greater than 91% accuracy, exhibiting successful separation of single-cells into chronological stages. Error bars indicate standard error. **(c)** Balloonplots showing the age-classification of de-novo sequenced beta-cells. GERAS predicted the age of the cells from independent sources with greater than 92% accuracy, showcasing the robustness of the model in handling biological and technical noise. **(d)** The capacity of GERAS to perform regression analysis was tested using cells with ages in-between the chronological stages used to train GERAS. More than 97% of the cells from the intermediate time-points classify in the nearest-neighbor stages. Number of cells for each condition is denoted by ‘n’.

For training GERAS, 80% of the beta-cells were randomly chosen. Optimal normalizing hyperparameters determined by cross-validation were used for training the final predictor. Following development, we estimated the contribution of the 1000 input genes towards accurate predictions (Supplementary Fig. S4, Supplementary Table S1). The estimation showed that the input genes displayed a wide distribution of importance towards the accuracy of prediction. Notably, some of these genes were previously implicated in diabetes (Supplementary Fig. S4b). Using the trained GERAS, internal validation was carried out with a test set comprising the remaining 20% of the cells from each chronological stage. The cells of the test set had never been shown to GERAS. Internal validation achieved an overall accuracy (proportion of cells for which the predicted stage matched the real stage) of 91% (Fig. 1b). This demonstrates the success of GERAS in classifying individual cells into chronological stages based solely on their mRNA expression profile.

Next, we wanted to understand the robustness of GERAS under biological and technical noise, typically encountered in batch measurements of single-cells. To this end, we performed external validation using independently sequenced beta-cells. We sequenced a new batch of beta-cells from adolescent animals (4 mpf) and used GERAS to predict their chronological age. All cells from this independent cohort were classified as ‘Adolescent’ (100% accuracy), the ground truth for the stage of the cells (Fig. 1c). Additionally, we tested the performance of GERAS with beta-cells sequenced using alternative pipelines.

Specifically, we utilized the C1-Chip platform from Fluidigm to sequence a new batch of beta-cells from adolescent animals (3 mpf). GERAS achieved 92.3% success in correctly classifying the cells from the new batch as ‘Adolescent’ (Fig. 1c). These data underscore the potential of GERAS in effectively handling batch effects.

To test the performance of GERAS on a regression task, we evaluated the model’s ability to classify cells obtained from time-points in-between the discrete chronological stages we used for training. For interpolation, we collected beta-cells from animals aged 1.5 mpf (juvenile) or 9 mpf (adult) since these ages were not part of the model’s constituent stages. GERAS classified 50% of the beta-cells from 1.5 mpf animals as ‘Juvenile’, and 47.3% as ‘Adolescent’ (Fig. 1d). Thus, GERAS classified 97.3% of beta-cells in time-periods neighboring the actual age of the sample. Similarly, 31% of the beta-cells from 9 mpf animals were classified as ‘Adolescent’, and 69% as ‘Adult’ (Fig. 1d). None (0%) of the cells were attributed to the ‘Juvenile’ stage, further strengthening the interpolation capacity of GERAS. Taken together, these results demonstrate that our model divides the continuous time variable into discrete but linearly-ordered stages, thereby allowing regression analysis of the data.

### GERAS evaluates the impact of an environmental factor on cellular age

The rate of aging is susceptible to modifications ^8^ and nutritional cues have been noted to alter aging in many organisms ^9,10^. To investigate the effect of altering nutritional cues on cellular age, we employed the ability of GERAS to handle batch effects and interpolation. Specifically, we focused on studying the impact of calorie intake on beta-cell aging. We separated 3 mpf adolescent zebrafish siblings into two groups. One group was fed three times a day with *Artemia*, a typical fish diet consisting of living prey with a relatively high amount of fat and carbohydrates ^43^. The other group was placed on intermittent feeding with normal feeding performed on alternate days (Fig. 2a). After one month, the beta-cells were isolated and the age of individual beta-cells was evaluated using GERAS for each group. The analysis showed a striking difference in age between the two sets of beta-cells obtained from coeval adolescent zebrafish (Fig. 2a). While 65% of the beta-cells from zebrafish on intermittent feeding were classified as ‘Adolescent’, only 23% of the beta-cells from three-times-a-day-fed animals were similarly classified; the rest 77% were categorized as ‘Adult’. This difference in classification of the beta-cells isolated from animals of the same age suggests that higher-caloric intake expedites the aging of young beta-cells. Moreover, it shows the utility of GERAS in evaluating a pro-aging factor.

**Figure 2:**
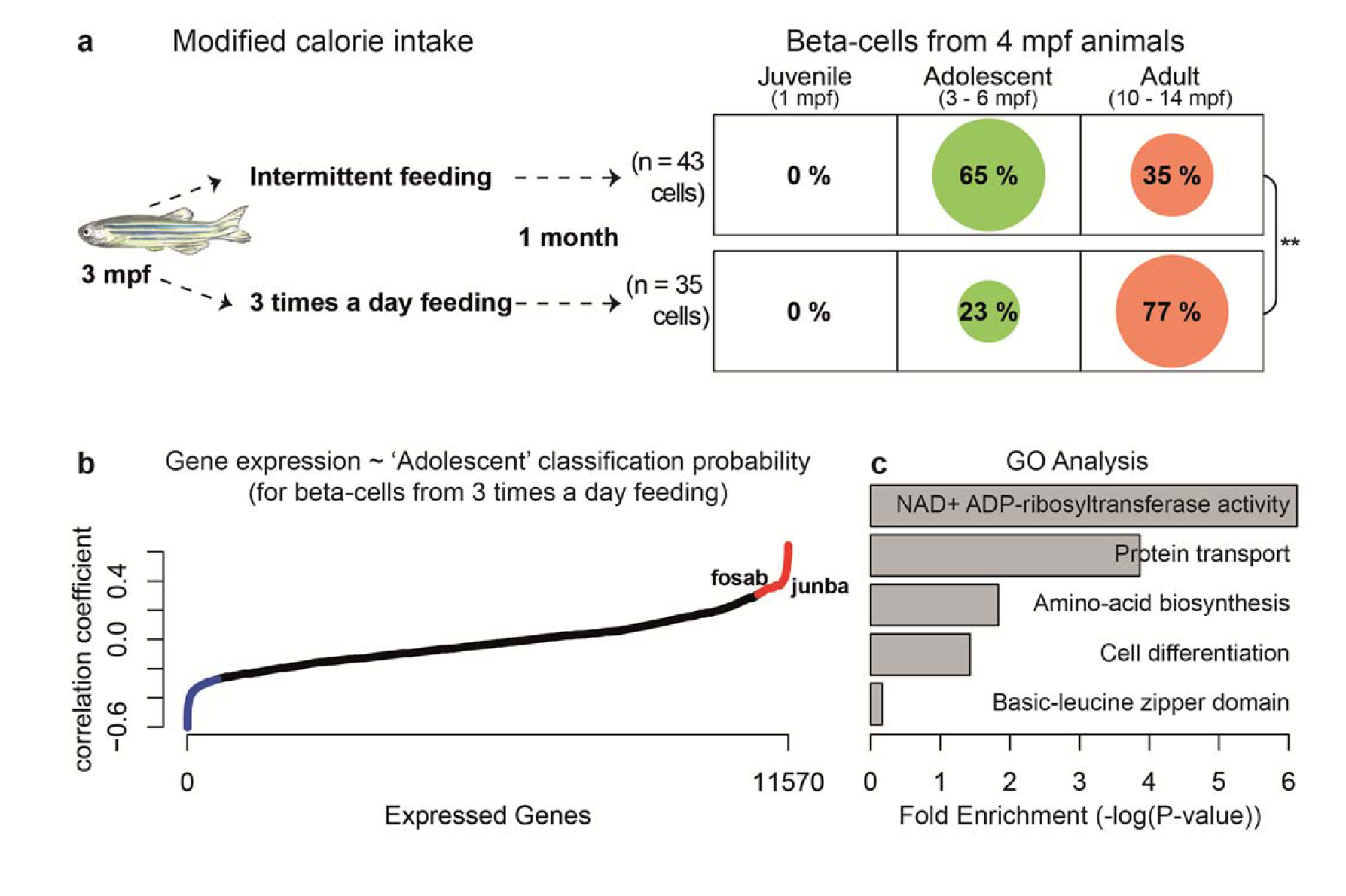
Impact of calorie intake on the chronological stage of zebrafish beta-cells. **(a)** The impact of calorie intake on the predicted age of beta-cells was investigated. Statistically, a higher proportion of beta-cells from 4 mpf animals fed three-times-a-day classified as ‘Adult’, as compared to cells from animals on intermittent feeding, in which a majority of the cells (67%) classified as adolescent. (Fisher’s Exact Test, **p-value < 0.01). **(b)** To identify the genes contributing to chronological stage classification, correlation analysis was performed. To this end, all beta-cells from the group fed three-times-a-day were used to calculate the correlation coefficient between gene expression and the probability of the cell to be classified in the ‘Adolescent’ stage. The Y-axis denotes the correlation coefficient and the X-axis depicts all the genes expressed in the beta-cells. The extreme fifth-percentile values are colored, with the red marking the top 5^th^ percentile (positive correlation) and blue marking the bottom 5^th^ percentile (negative correlation). Genes with positive correlation, which include *junba* and *fosab*, contribute towards classification in the ‘Adolescent’ stage as opposed to classification in the ‘Adult’ stage, thereby increasing the probability of a cell being classified as younger. **(c)** Gene-ontology (GO) analysis using DAVID ^44^ for genes in the extreme fifth-percentile. This analysis includes the genes exhibiting negative (blue in b) and positive (red in b) correlation. Zebrafish illustration provided with permission.

### GERAS-based predictions lead to discovery of a molecular factor involved in aging

To identify molecular players underlying the accelerated aging of beta-cells with higher-calorie intake, we harnessed the heterogeneity in the chronological stage predictions along with the inherent heterogeneity in gene expression within single cells. In our framework, chronological stage predictions can be easily converted to classification probability by using the output of ‘softmax’ layer (Fig. 1a and Methods). This transforms discrete classifications into a continuous probability distribution (Supplementary Fig. S5). Taking advantage of this approach, we calculated the correlation between the probabilities of the beta-cells to be classified in the younger (‘Adolescent’) stage with the mRNA expression levels of all 11,570 genes expressed in the beta-cells (Supplementary Fig. S5). For correlation analysis, genes with positive correlation increase the chance of the cell being classified in the younger stage, while a negative correlation enhances the chance of classification in the older stage. The correlation analysis for beta-cells from three-times-a-day fed animals revealed 1158 genes exhibiting high (positive or negative) correlation with predictive probability (Fig. 2b, Supplementary Table S2 and S3). Unbiased gene ontology analysis using DAVID ^44^ revealed involvement of the highly correlated genes in aging-related pathways, including cellular differentiation, protein transport ^45,46^, amino acid biosynthesis ^47,48^, NAD+ ADP-ribosyltransferase activity ^49^ and basic-leucine zipper domain containing transcription factors ^50^ (Fig. 2c). In particular, there was a positive correlation with the transcription factors *junba* and *fosab*, suggesting a role for these genes in the classification of the beta-cells to the younger, ‘Adolsecent’, stage (Fig. 2b). Additionally, in our primary mRNA expression data of beta-cells from three chronological stages, *junba* and *fosab* displayed significant down-regulation with age (Supplementary Fig. S6). Notably, *junba*, was not one of the 1000-input genes utilized by GERAS for generating predictions, demonstrating the capacity of correlation analysis to identify genome-wide candidate genes.

Based on the observation that *junba* expression in beta-cells declines with age, and its positive correlation with the classification of beta-cells from animals on a higher-calorie diet to the younger stage, we decided to investigate the biological impact of reducing *junba* function. For this, we overexpressed a dominant negative version of *junba* specifically in beta-cells (using an *ins:nls-BFP-2A-DN-junba* construct) (Supplementary Fig. S7a). The expression of *nls-BFP-2A-DN-junba* was induced in the background of the beta-cell specific fluorescence ubiquitination cell cycle indicator (FUCCI)-reporters ^51,52^, allowing identification of beta-cell’s cell-cycle stage (Supplementary Fig. S7b, c). Comparison between the juxtaposed *DN-junba*-expressing and control cells within islets from juveniles (1 mpf), a stage associated with high rates of beta-cell proliferation ^51^, showed a 50% decline in proliferation upon *DN-junba* expression (Fig. 3a, b). Thus, blocking *junba* function can reduce the proliferation of juvenile beta-cells. Since the reduction in proliferation of beta-cells is a hallmark of aging ^41^, our results suggest that declining *junba* expression might underlie this reduction.

**Figure 3:**
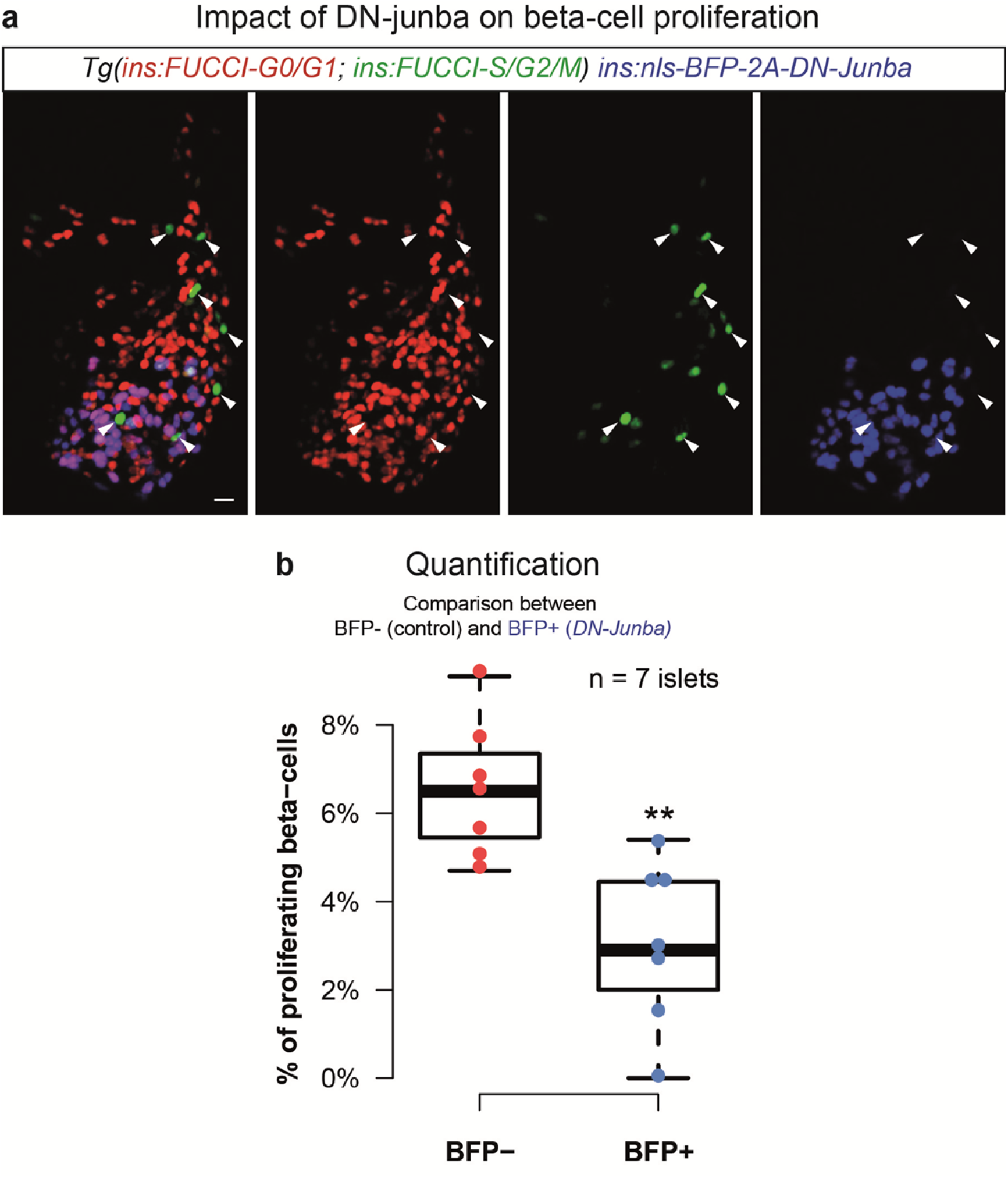
Inhibition of *junba* reduces the proliferation of zebrafish beta-cells. **(a)** Maximum intensity confocal projections of islet from 30 dpf animal showing mosaic expression of *nls-BFP-2A-DN-junba* (blue) together with *Tg(ins:FUCCI-S/G2/M)* (green) and *Tg(ins:FUCCI-G0/G1)* (red). Arrowheads mark proliferating beta-cells, as indicated by the presence of green fluorescence and absence of red fluorescence. Scale bar 10 μm. **(b)** Tukey-style boxplots showing the percentage of proliferating beta-cells among BFP+ and BFP-cells. BFP+ cells co-express *DN-junba*, while the BFP-cells act as internal control. The BFP+ cells show a statistically significant decrease in the proportion of proliferating cells (t-test, **p-value <0.01). ‘n’ denotes number of islets.

### A single model for chronological stage classification of the entire human pancreatic cells

Next, to test the applicability of our framework beyond the scope of zebrafish beta-cells, we developed a classifier for human cells using the entire ensemble of pancreatic cells. The pancreas, a gland located in the abdomen, is involved in metabolic regulation and food digestion. Metabolic regulation is accomplished by the endocrine part of the pancreas, which chiefly consists of beta-, alpha-, and delta-cells. Food digestion, on the other hand, is contributed by the exocrine part of the pancreas, composed of ductal and acinar cells. An important characteristic of pancreatic cells is the presence of cell-specific marker genes, allowing computational segregation of the various cell-types based on mRNA expression levels (Methods). To develop the classifier for human pancreatic cells, we obtained singlecell mRNA expression profiles from Enge et al. Their study generated single-cell transcriptomes from pancreatic cells of eight healthy individuals belonging to three discrete stages ^27^: Juvenile (1 month, 5 and 6 years), Young (21 and 22 years), and Middle (38, 44 and 54 years) (Fig. 4a). Without segregating the data by cell-type, we trained GERAS to predict the chronological stage for the entire ensemble of pancreatic cells. The trained GERAS, utilizing inputs from multiple genes (Supplementary Fig. S8, Supplementary Table S4), achieved an overall accuracy of 95% on the test set (Fig. 4b). Upon segregating the results by cell type, based on the expression of their respective markers, we found that GERAS displayed >90% accuracy for each major cell-type of the pancreas (Fig. 4b’), demonstrating the feasibility of developing a single age classifier for the multiple cell types of the pancreas.

**Figure 4:**
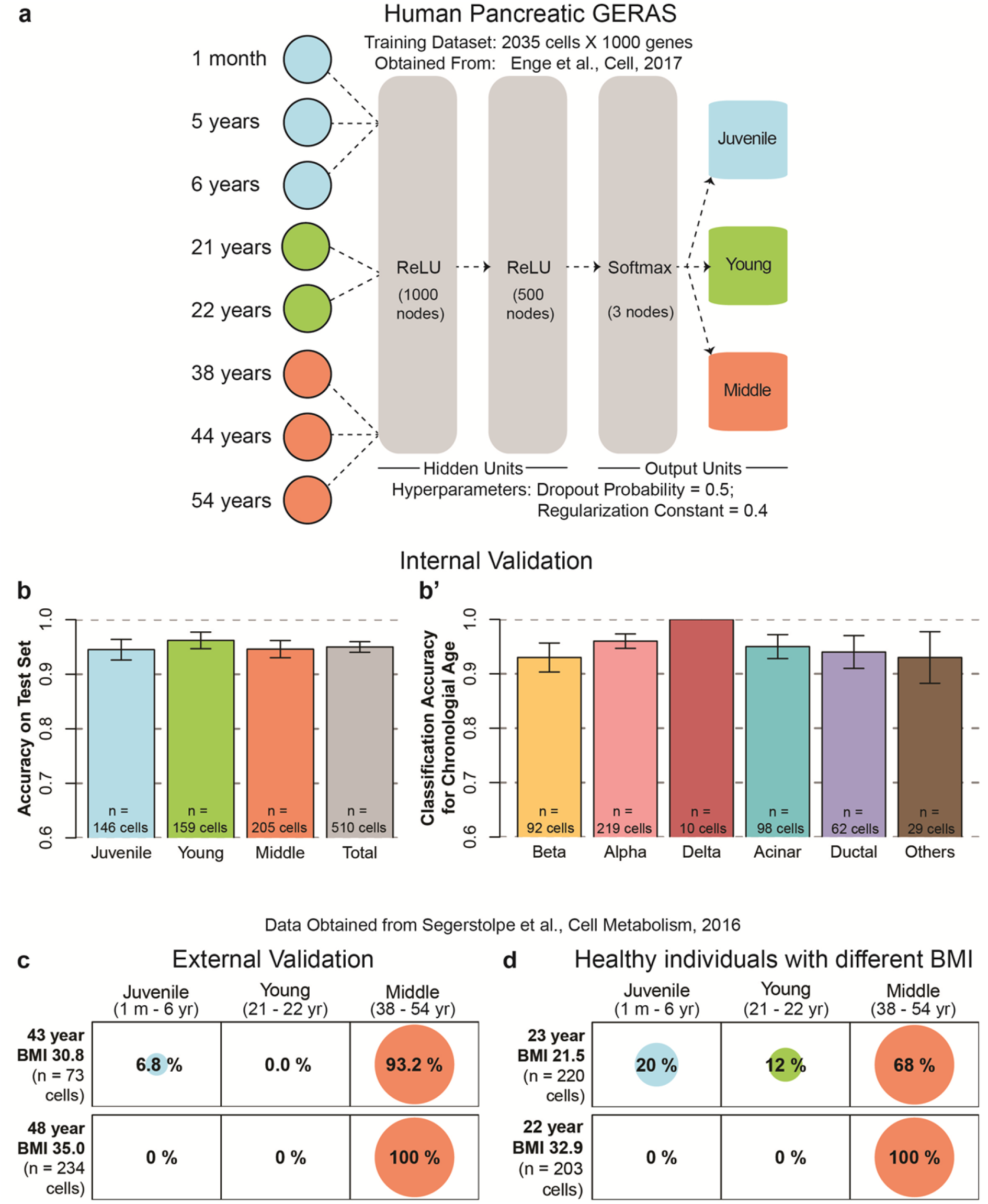
A Chronological age classifier for human pancreatic cells. **(a)** A single chronological age classifier for the entire ensemble of human pancreatic cells using machine learning. No cell-type segregation was performed during training. **(b)** Barplot showing the accuracy of GERAS on classifying the age of pancreatic cells that were not used for training the model. An accuracy of 95% was achieved for cells previously unseen by GERAS. (**b’**) The classification accuracy of GERAS on the previously unseen pancreatic cells after segregating them into major cell-types. Classification accuracy equals the proportion of cells for which the predicted stage matched the actual stage. For each cell-type, greater than 93% accuracy was achieved. Error bars indicate standard error. **(c)** External validation for the classifier was provided by human pancreatic single-cell mRNA expression data obtained from an independent publication. Cells from individuals belonging to the ‘Middle’ (38 - 54 years) stage of the classifier displayed greater than 93% accuracy. **(d)** Balloonplot showing classification of cells from individuals with similar chronological age but different BMI. In individuals with normal BMI, 32% of the cells were classified in ‘Juvenile’ and ‘Young’ stages, while none (0%) of the cells from individuals with obese BMI were similarly classified. Number of cells for each condition is denoted by ‘n’.

As an additional validation, a second assessment with human cells was undertaken by utilizing the single-cell mRNA expression profiles of human pancreatic cells from a publication by Segerstolpe et al. ^53^. This independent cohort contains single-cell transcriptomes from pancreata of six healthy individuals ranging from 22 - 48 year of age. Additionally, the body mass index (BMI) for each individual was reported, allowing comparisons between individuals with similar chronological age but different body weight. Using GERAS trained with the human data from Enge et al., we predicted the chronological stage of the cells from two individuals (aged 43 and 48 years) belonging to the ‘Middle’ age group (38 - 54 years). The predictions displayed >93% classification accuracy (Fig. 4c). This high accuracy of prediction on data from a second independent source further strengthens the external validation of our model. Next, we utilized the data from two individuals, aged 23 and 22 years. Despite the proximity in their chronological age, these two individuals differed in their BMI values (21.5 - normal and 32.9 - obese, respectively). Strikingly, our analysis revealed different classification pattern for data from each of these individuals: while 32% of the cells from the 23 year old with normal BMI were classified in the younger stages, none of the cells from the 22 year old with obese BMI fell in similar stages (Fig. 4d). Following this observation, we calculated the classification probability of the all six individuals in relation to their BMI. The probability results from our analysis suggest that an obese BMI correlates with an increased probability for the cells to be classified in an older stage (Supplementary Fig. S9). We recommend exercising caution while interpreting this result due to the multiple confounding factors associated with human samples that we could not control for. A GERAS developed with cells from individuals encompassing a wider distribution of age and BMI range would be desired for stronger conclusions. Nevertheless, the successful age classification of an entire human organ and its external validation, demonstrate the adaptability of our framework to diverse cell-types, thereby establishing the universality of the approach.

## DISCUSSION

In this study, we have presented a method that provides the blueprint for developing predictive classifier for cellular aging. Our chronological stage predictor efficiently handles biological and technical noise, and functions robustly on a diverse cell population. The temporal classifier was developed in an unbiased, data-driven manner. Genes for building the predictor were not selected based on their differential expression with time. The classifier predicted the chronological age solely from the expression profile of the top 1000 most variable genes. The algorithm, however, did not use all genes uniformly. Instead, varying levels of importance were attributed to the input genes (Supplementary Fig. S4, S8). Multiple genes exhibiting high importance for successful classification show an existing association with metabolic and age-related degenerative disorders. For instance, the human pancreatic GERAS ascribes high importance to Amyloid precursor protein (APP), which is associated with Alzheimer’s disease, and also recently implicated in pancreatic biology ^54^ In the future, it would be worthwhile to test the biological functions for the genes selected by the classifier, and to follow-up on them as potential biomarkers of the aging process.

The predictive power of the framework is not restricted to classification tasks. The discrete classifications can be readily converted to a continuous probability distribution (Supplementary Fig. S5). This characteristic can be exploited to shed light on the molecular factors controlling the rate of aging. We used this feature on beta-cells displaying accelerated aging in response to a higher calorie diet (Fig. 2a). Correlating the probability distribution with gene expression enabled identification of candidate genes involved in the aging process (Fig. 2b, c). Such analysis was possible due to the single-cell-centric nature of our approach, and would be missed out with bulk sequencing in which the cellular variability is averaged out. Follow-up analysis using a genetic technique (Supplementary Fig. S7) verified the role of one candidate gene, *junba*, in regulating the proliferation of beta-cells (Fig. 3). It is important to note that the mosaic analysis was performed in whole islets without any tissue dissociation, thus avoiding any dissociation-specific modification in cell physiology ^55^. However, a reduction in proliferation represents one aspect of the aging process, and additional roles for *junba* activity during the aging process still need verification. Nonetheless, the age-dependent reduction of *Junb*, the mammalian homologue of *junba*, has been implicated in post-natal maturation of mouse beta-cells ^56^. It would be of interest to follow-up on these results and study the connection between aging and *Junb* activity in mammalian models.

Importantly, beta-cells from animals fed three-times-a-day revealed a diversity in their classification. Notably, 23% of the beta-cells were classified in the younger stage, suggesting cellular heterogeneity in the aging process. This was additionally observed during the interpolation analysis (Fig. 1d), in which cells from intermediate time-points classified in the two adjacent stages. Asynchronous cellular aging in beta-cells was recently hypothesized using histological analysis ^57^ Quantifying the extent of heterogeneity in the aging process while capturing the mRNA expression profile, made possible by our framework, provides an exciting opportunity for understanding the molecular underpinnings of heterogeneous cellular aging.

Our machine-learning based framework has high flexibility in its design and execution, which can be exploited to develop predictive models based on diverse biological parameters. Moreover, the inputs to the predictor are not limited to mRNA expression levels but can be extended to include other covariates. With improvements in single cell epigenetics^58^, new models integrating both genetic and epigenetic changes could be built to improve accuracy and resolution.

Our framework is based on the assumption that chronological age provides a useful metric for the modeling of age. Chronological age is an easily observable fact, and this provided the ground truth for training and testing our models. The aging trajectory provided by the use of chronological age served as benchmark for all predictions generated by the framework. However, chronological age does not always correlate well with development of disease and mortality ^59^ Previous studies have introduced the concept of biological age ^60,61^, a metric that correlates better than chronological age with pathological conditions. However, the determination of biological age requires training, testing and verification of regression models. This leads to the biological age being defined as per the computation model, which can result in very low overlap between different measures of biological age ^62^ In the future, it would be worthwhile to generate two-tier models combining the information from models based on chronological and biological age.

We developed our model with the idea in mind to be able to detect premature aging. However, individual responses might differ towards the factors that lead to accelerated aging. For instance, within the population of humans with an obese BMI, the ‘metabolically healthy obese’ group exhibits lower risk for complications as compared to the ‘metabolically unhealthy obese’ ^63,64^. Further work needs to be done to identify individual risk-factors associated with premature aging. This would be necessary for recommendations of preventive therapies.

The predictors presented in this study are restricted by sequencing platforms and the specific tissues utilized for training them. This limits their immediate adaptation. The predictors are built with data generated from Smart-Seq2 sequencing pipeline, which captures the full-length mRNAs with high transcriptome coverage. The predictor might be unable to handle the data from Drop-seq or MARS-seq, protocols that sequence the 3’-end of mRNA and provide lower-coverage ^42^ Computational efforts for eliminating the idiosyncrasies of individual platforms ^65^ would help to remove this restriction. Additionally, the predictors do not extend beyond the currently described tissues. Investigators interested in the aging of other cells, for instance muscle, would need to develop and validate *de-novo* predictive models. Nevertheless, we expect the groundwork presented here to help with the development of predictive models. Further improvements of our approach could expedite the identification of age-modifying factors, which are important regulators of development and disease.

## CONCLUSION

Here we developed a machine learning based platform that successfully predicts the chronological stage of individual cells. We show the framework’s robustness in handling multiple sample processing pipelines, time-points that fall between the discrete chronological stages, and diversity in cell types. The framework’s capability to characterize aging factors was demonstrated through evaluation of the impact of a higher-calorie feeding on beta-cell aging. The predictive power of the framework was further harnessed to discover *junba* as a candidate gene that maintains the proliferative beta-cell state, a characteristic trait of younger beta-cells. Broad applicability of the framework was demonstrated by predictions on the entire human pancreatic tissue. We anticipate that the robustness and flexibility exhibited here will enable the development of aging models for multiple tissues, opening the possibility of detecting premature aging and preventing pathological developments. To maximize the accessibility and impact of the study, the framework is openly shared on github ^66^, and a user-friendly, graphical interface is provided for generating predictions from trained models.

## METHODS

### Zebrafish strains and husbandry

Wild-type or transgenic zebrafish of the outbred AB, WIK or a hybrid WIK/AB strain were used in all experiments. Zebrafish were raised under standard conditions at 28°C. Animals were chosen at random for all experiments. Published transgenic strains used in this study were *Tg(ins:BB1.0L; cryaa:RFP)*^36^; *Tg(ins:FUCCI-G1)*^s948^ ^51^; *Tg(ins:FUCCI-S/G2/M)*^s946^ ^51^. Experiments were conducted in accordance with the Animal Welfare Act and with permission of the Landesdirektion Sachsen, Germany (permits AZ 24-9168, TV38/2015, T12/2016, and T13/2017).

### Single cell isolation of zebrafish beta-cells

Primary islets from *Tg(ins:BB1.0L; cryaa:RFP)* zebrafish were dissociated into single cells and sorted using FACS-Aria II (BD Bioscience). Islets were dissociated into single cells by incubation in TrypLE (ThermoFisher, 12563029) with 0.1% Pluronic F-68 (ThermoFisher, 24040032) at 37 °C in a benchtop shaker set at 450 rpm for 30 min. Following dissociation, TrypLE was inactivated with 10% FBS, and the cells pelleted by centrifugation at 500g for 10 min at 4 °C. The supernatant was carefully discarded and the pellet re-suspended in 500 uL of HBSS (without Ca, Mg) + 0.1% Pluronic F-68. To remove debris, the solution was passed over a 30 μm cell filter (Miltenyi Biotec, 130-041-407). To remove dead cells, calcein violet (ThermoFisher, C34858) was added at a final concentration of 1 μM and the cell suspension incubated at room temperature for 20 minutes. The single cell preparation was sorted with the appropriate gate for identification of beta-cells (RFP+ and calcein+) (Supplementary Fig. S1). FACS was performed through 100 μm nozzle with index sorting.

### Single cell mRNA sequencing of zebrafish beta-cells from 96-well plates

Cells were sorted into a 96-well plate containing 2 μl of nuclease free water with 0.2% Triton-X 100 and 4 U murine RNase Inhibitor (NEB), spun down and frozen at −80°C. After thawing the samples, 2 μl of a primer mix was added (5 mM dNTP (Invitrogen), 0.5 μM dT-primer*, 4 U RNase Inhibitor (NEB)). RNA was denatured for 3 minutes at 72°C and the reverse transcription was performed at 42°C for 90 min after filling up to 10 μl with RT buffer mix for a final concentration of 1x superscript II buffer (Invitrogen), 1 M betaine, 5 mM DTT, 6 mM MgCl2, 1 μM TSO-primer*, 9 U RNase Inhibitor and 90 U Superscript II. After synthesis, the reverse transcriptase was inactivated at 70°C for 15 min. The cDNA was amplified using Kapa HiFi HotStart Readymix (Peqlab) at a final 1x concentration and 0.1 μM UP primer under following cycling conditions: initial denaturation at 98°C for 3 min, 22 cycles [98°C 20 sec, 67°C 15 sec, 72°C 6 min] and final elongation at 72°C for 5 min. The amplified cDNA was purified using 1x volume of hydrophobic Sera-Mag SpeedBeads (GE Healthcare) and DNA was eluted in 12 μl nuclease free water. The concentration of the samples was measured with a Tecan plate reader Infinite 200 pro in 384 well black flat bottom low volume plates (Corning) using AccuBlue Broad range chemistry (Biotium).

For library preparation, 700 pg cDNA in 2 μl was mixed with 0.5 μl tagmentation enzyme and 2.5 μl Tagment DNA Buffer (Nextera DNA Library Preparation Kit; Illumina) and tagmented at 55°C for 5 min. Subsequently, Illumina indices were added during PCR (72°C 3 min, 98°C 30 sec, 12 cycles [98°C 10 sec, 63°C 20 sec, 72°C 1 min], 72°C 5 min) with 1x concentrated KAPA Hifi HotStart Ready Mix and 0.7 μM dual indexing primers. After PCR, libraries were quantified with AccuBlue Broad range chemistry, equimolarly pooled and purified twice with 1x volume Sera-Mag SpeedBeads. This was followed by Illumina sequencing on a Nextseq500 aiming at an average sequencing depth of 0.5 million reads per cell.

* dT primer: Aminolinker-AAGCAGTGGTATCAACGCAGAGTCGAC T(30) VN
* TSO primer: AAGCAGTGGTATCAACGCAGAGTACATggg
* UP primer: AAGCAGTGGTATCAACGCAGAGT

### Single cell mRNA sequencing of zebrafish beta-cells with the C1 system

The <u>C1™ Single-Cell mRNA Seq 10-17 μm</u> IFC (© Fluidigm Corporation, CA, USA) was used to perform mRNA sequencing on single cells. In general, the protocol (PN 100-7168 L1) suggested by the manufacturer was followed, with some modifications. 1200 cells in PBS were directly sorted by FACS into the inlet, mixed 3:2 with suspension reagent, resulting in a final volume of 6 μl. Cells were loaded with the mRNAseq: Cell load protocol, without staining on the IFC. For RT and amplification, the mRNA Seq: RT & Amp script was run with the following cycling parameters: 1x 98°C 1 min, 5x (95°C 20-45 sec, 59-49ºC with 0. 3°C increment/cycle 4 min, 68°C 6 min) 9x (95°C 20-45 sec, 65-49ºC with 0.3°C increment/cycle 30 sec, 68°C 6 min) 7x (95°C 30-45 sec, 65-49°C with 0.3°C increment/cycle 30 sec, 68°C 7 min) and 72°C 10 min using SMART-Seq v4 Ultra Low Input RNA Kit for Sequencing (Takara BIO USA, INC.). For library preparation, 2 μl cDNA were mixed with 0. 5 μl tagmentation enzyme and 2.5 μl Tagment DNA Buffer (Nextera DNA Library Preparation Kit; Illumina) and tagmented at 55°C for 5 min. Illumina indices were added by PCR with the following cycling conditions: 1x (72°C 3 min, 98°C 30 sec), 12 x (98°C 10 sec, 63°C 20 sec, 72°C 1 min), 1x (72°C 5 min), using KAPA Hifi HotStart Ready Mix and 0.7 μM final dual indexing primers. Libraries were quantified, equimolarly pooled and purified twice with 1x volume Sera-Mag SpeedBeads. Illumina sequencing (75bp SE) was done on a Nextseq500 aiming to achieve an average sequencing depth of 0.5 million reads per cell.

### Mapping of read counts and quality control

Raw reads in fastq format were trimmed using trim-galore with default parameters to remove adapter sequences. Trimmed reads were aligned to the zebrafish genome, GRCz10, using HISAT2 ^67^ with default parameters. htseq-count ^68^ was used to assign reads to exons thus eventually getting counts per gene. Using cells that were utilized for developing zebrafish GERAS (see next section), the following quality control parameters were obtained (Supplementary Fig. S2):

1. The median and median absolute deviation (MAD) for total reads
2. The median and MAD for % of mitochondrial reads
3. The median and MAD for % spike-ins
4. Number of detectable genes

Cells passed quality control if they belonged to median ± 3*MAD bracket for 1-3 and contained more than 1500 genes. Read counts for all cells that passed quality control are available at: https://sharing.crt-dresden.de/index.php/s/zcQ14AMGJAevokU.

### Pseudotemporal ordering of zebrafish beta-cells

Unsupervised pseudotemporal ordering of zebrafish beta-cells was carried out using the read counts from beta-cells isolated from seven different ages. The cells were grouped in three stages before analysis: ‘Juvenile’ (1 mpf), ‘Adolescent’ (3, 4, 6 mpf) and ‘Young’ (10, 12, 14 mpf). Ordering was carried out using Monocle ^16^, as outlined in the vignette for Monocle2. The analysis is shared online as Monocle.R.

### Development of GERAS for zebrafish beta-cells

For development of GERAS for zebrafish beta-cells, read counts were used from seven ages of zebrafish: 1 mpf, 3 mpf, 4 mpf, 6 mpf, 10 mpf, 12 mpf and 14 mpf. The 3 mpf and 6 mpf stages contained two batches of beta-cells collected and sequenced on different days. Each batch of cells originated from six zebrafish. Read counts were normalized to transcripts per million (TPM) using the formula:

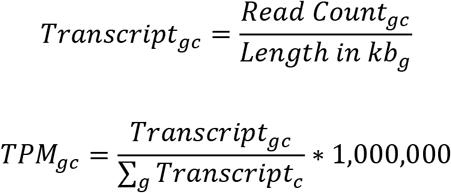

where for gene *g* and cell *c*, *Transcript_gc_* are the number of transcripts calculated by dividing the read counts to the length of the gene in kb, and TPM is the proportion of the gene’s transcripts among per million of total cellular transcripts.

The entire dataset containing 508 beta-cells were randomly divided into 80%-20% train-test set. Genes were sorted in descending order according to their expression variability (calculated by ‘median absolute deviation’) in the entire dataset. The top 1000 most variable genes were used for developing a four-layer fully connected neural network (Fig. 1a). The neural network contained two hidden layers with rectified linear unit (ReLU) activation function, and a softmax output layer. The network was trained to classify the pancreatic cells into three chronological ages: Juvenile (1 month post-fertilization (mpf)), Adolescent (3, 4 and 6 mpf) and Adult (10, 12 and 14 mpf). During training, a five-fold cross-validation was repeated three times over a grid of values for regularization hyperparameters: dropout frequency (0.4 to 0.9 in steps of 0.1) and regularization constant (0.4 to 1.6 in steps of 0.2). The combination with the highest cross-validation accuracy was taken as the optimal value, and a final model was trained using the entire training set and the optimal regularization hyperparameters. The entire network was implemented in R using TensorFlow API. An Rmarkdown report detailing the development of zebrafish beta-cell GERAS is available at https://github.com/sumeetpalsingh/GERAS2017/blob/master/GERAS_Tf_Zf.html^66^.

The trained model was used to predict the chronological age of the test set. Accuracy was calculated as the proportion of cells for which the prediction matched the chronological age. By considering each prediction as a binomial distribution (a ‘Juvenile’ cell can be classified as ‘Juvenile’ or ‘Not Juvenile’), the standard error was calculated using the following formula:

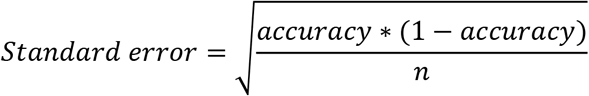

where *n* is the number of cells tested.

### Prediction of chronological age using GERAS for zebrafish beta-cells

For external validation (4 mpf and 3 mpf C1-sample) and interpolation (1.5 mpf and 9 mpf), new batches of zebrafish beta-cells were isolated in 96-well plates and sequenced. Quality controlled raw counts were obtained as outlined above. The raw counts were normalized to TPM values, which were then used to predict the chronological stage using pretrained GERAS. Results were depicted as balloonplots, where a grid contains dots whose size reflects the percentage of cells classified in the corresponding group.

### Assessing the impact of calories on the chronological age of zebrafish beta-cells using GERAS

Twelve zebrafish at 3 mpf from the same clutch were separated into two groups of 6 animals each. Both groups were fed with their normal feed of freshly hatched *Artemia* (brine shrimp). The intermittent feeding group was fed on alternate day, while the other group was fed three times daily with intervals of at least two hours between the feedings. Amount of food eaten by each animal was not controlled. After a month, the beta-cells were isolated into 96-well plates using FACS. The cells were processed and sequenced together. TPM-normalized counts from the cells were used to predict the chronological age using GERAS.

### Correlation analysis and gene ontology (GO) analysis

Correlation analysis was carried out for beta-cells collected from the three-times-a-day animals. These beta-cells classified in ‘Adolescent’ and ‘Adult’ stage (Fig. 2a). The analysis calculated the correlation between the probability of a cell to be classified in the younger (‘Adolescent’) stage and the mRNA expression of genes. To obtain the classification probability, the softmax for the ‘Adolescent’ stage was calculated from the output layer of GERAS (Fig. S5). For this, a function (model_softmax) was written that takes the log2-transformed normalized values of single cells, performs forward propagation through GERAS till the softmax layer, and returns the output. The output contains the probability for the particular cell to classify in all the three stages (‘Juvenile’, ‘Adolescent’, and ‘Adult’). The function is deposited as source/model_softmax.R ^66^. The probability for ‘Adolescent’ stage was extracted from this output.

Correlation coefficient was calculating using the cor(classification probability, gene expression) function in R. The calculation was restricted to genes expressed in more than 10% of the cells (11,570 genes). This gave a correlation value for each gene expressed in beta-cells from three-times-a-day animals. The values were sorted in ascending order and plotted in Fig. 2b. The genes with the highest positive correlation were identified as the top fifth-percentile, and the genes with the highest negative correlation were identified as the lowest fifth-percentile. These genes were further used for unbiased gene ontology (GO) analysis using DAVID ^44^ As background for GO analysis, the list of expressed genes was used.

### Construction of the *ins:nls-BFP-T2A-DN-junba; cryaa:RFP* plasmid

To generate *ins: nls-BFP-T2A-DN-junba;cryaa:RFP*, a vector was created by inserting multiple cloning sites (MCS) downstream of the insulin promoter to yield *ins:MCS; cryaa:RFP*. To do so, the plasmid *ins:mAG-zGeminin;cryaa:RFP* was digested with EcoRI/PacI and ligated with dsDNA generated by annealing two primers harboring the sites EcoRV, NheI, NsiI, SalI and flanked by EcoRI/PacI overhangs. The plasmid pUC-Kan consisting of the DN-junba (junba^157-325^, consisting of only the DNA binding domain^69^) fused to *nls-BFP* via T2A sequence flanked by EcoRI/PacI sites was synthesized from GenScript. *ins:MCS;cryaa:RFP* and the plasmid *pUC-nls-BFP-T2A-DN-junba* were subsequently digested with EcoRI/PacI to yield compatible fragments, which were ligated together to yield the final construct. The entire construct was flanked with I-SceI sites to facilitate genomic insertion.

### Analysis of proliferation using mosaic expression of *DN-junba*

To identify proliferating beta-cells, the zebrafish beta-cell specific FUCCI system^51^ was used by crossing *Tg(ins:FUCCI-G1)* with *Tg(ins:FUCCI-S/G2/M)*. Embryos obtained from the mating were injected with *ins:nls-BFP-T2A-DN-junba;cryaa:RFP* plasmid, along with I-SceI, to facilitate mosaic integration into the genome. At 30 dpf, animals were euthanized in Tricaine and dissected to isolate the islets. The isolated islets were fixed in 4% paraformaldehyde (PFA) for 48 hours at 4°C, washed multiple times in PBS and mounted on slides for confocal microscopy. Confocal images were used for cell-counting. All the *Tg(ins*:FUCCI-S/G2/M)-positive cells (green fluorescence only) were counted manually within the BFP-positive and BFP-negative clones. Using Imaris (Bitplane), the total number of BFP-positive and beta-cells were calculated in the entire islet. For this, the “spots” function was used after thresholding. For calculating percentages (%), the following calculations were used:

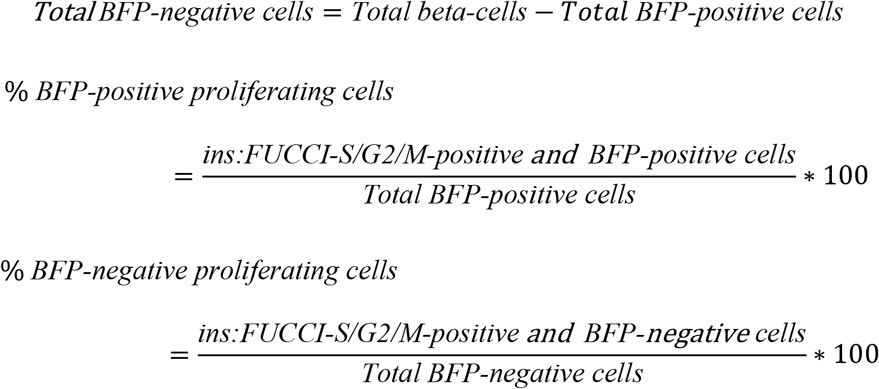

### Statistical analysis

Statistical analysis was performed using R. No animals were excluded from analysis. Blinding was not performed during analysis. Analysis of normal distribution was performed. To compare chronological age (Adolescent versus Adult) between beta-cell from intermittent feeding and three-times a day fed animals, Fisher’s exact test for count data (fisher.test(x = 2X2 matrix, alternative = "two.sided")) was performed. To compare the expression levels of *junba* and *fosab* between Juvenile, Adolescent and Adult, ANOVA followed by Tukey's range test (fit <-aov(Expression ∼ Stage); TukeyHSD(fit)) was performed. To compare the proliferation between *DN-junba* expressing cells with control cells, an unpaired two-tailed t-test with unequal variance (t.test (x = dataframe, alternative = "two.sided", paired = FALSE, var.equal = FALSE)) was used to calculate p-values. A p-value of less than 0.05 was considered statistically significant.

### Development of GERAS for human pancreatic cells

For development of GERAS for human pancreatic cells, read counts from Enge et al.^27^ were obtained from GEO: GSE81547. Read counts were normalized to reads per million (RPM) using the formula:

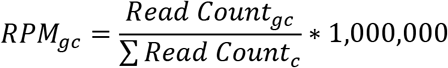

where for gene *g* and cell *c*, RPM_gc_ is the proportion of the gene’s reads among per million of the total cellular reads.

The entire dataset containing 2544 pancreatic cells was randomly divided into 80%-20% train-test set. Genes were sorted in descending order according to their expression variability (calculated by ‘median absolute deviation’) in the entire dataset. The top 1000 most variable genes were used for developing a four-layer fully connected neural network (Fig. 4a). The neural network contained two hidden layers with ReLU activation function, and a softmax output layer. The network was trained to classify the pancreatic cells into three chronological ages: Juvenile (1 month, 5 and 6 years), Young (21 and 22 years), and Middle (38, 44 and 54 years). During training, a five-fold cross-validation was repeated three times over a grid of values for regularization hyperparameters: dropout frequency (0.4 to 0.9 in steps of 0.1) and regularization constant (0.2 to 1.2 in steps of 0.2). The combination with the highest cross-validation accuracy was taken as the optimal value, and a final model was trained using the entire training set and the optimal regularization hyperparameters. The entire network was implemented in R using TensorFlow API. An Rmarkdown report detailing the development of human pancreatic GERAS is available at https://github.com/sumeetpalsingh/GERAS2017/blob/master/GERAS_Tf_Hs.html ^66^.

The trained model was used to predict the chronological age of the test set. Accuracy was calculated as the proportion of cells for which the prediction matched the chronological age. By considering each prediction as a binomial distribution (a ‘Middle’ cell can be classified as ‘Middle’ or ‘Not Middle’), the standard error was calculated using the following formula:

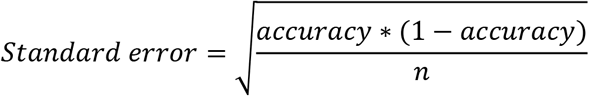

where *n* is the number of cells tested.

To calculate the accuracy and standard error per cell type, the expression levels of the following cell-specific markers were extracted for each cell: ‘INS’ (beta-cell), ‘GCG’ (alphacell), ‘SST’ (delta), ‘PRSS1’ (acinar) and ‘KRT19’ (ductal). A cell was classified if the expression value of any cell-specific marker exceeded 50 RPM, else it was classified as ‘Others’. For classification, the cell-type marker with the highest expression determined the cell type. Thus, a (theoretical) cell with RPM values of 1000 INS, 3 GCG, 4 SST, 0 PRSS1, 0 KRT19 was classified as beta-cell, while another (theoretical) cell with RPM values of 3 INS, 5 GCG, 7 SST, 1777 PRSS1, 9 KRT19 was classified as acinar cell. Cell-type specific cells present in the test set were used to calculate the accuracy per cell-type.

### Independent cohort of human pancreatic cells

For testing GERAS with external data, read counts of pancreatic single-cell data from Segerstolpe et al.^53^ were obtained from ArrayExpress (EBI) with accession number: <u>E-MTAB-5061</u>. The publication contained data from six healthy individuals. The entire data was stratified according to the individuals, and cells from each individual that passed quality-control according to Segerstolpe et al. were used for further analysis. Read counts from the cells were normalized to RPM for input to GERAS.

### Calculating classification probability for ‘Middle’ (38 - 54 years) stage

To calculate the probability that a particular cell would be classified to the ‘Middle’ stage, the softmax for the ‘Middle’ stage was calculated from the output layer of human pancreatic GERAS. For this, the function model_softmax was provided with the log2-transformed RPM values and used to calculate the probability for the particular cell to classify in all the three stages (‘Juvenile’, ‘Young’, and ‘Middle’). The probability for ‘Middle’ stage was extracted from this output.

### Prediction of chronological age using GERAS for human pancreatic cells

For predicting the chronological stage of cells belonging to individuals of age 22, 23, 43 and 48 years, RPM values from each individual were used as input to human pancreatic GERAS. Results were depicted as balloonplots, where a grid contains dots whose size reflects the percentage of cells classified in the corresponding group.

### Calculating variable importance for GERAS

Variable importance was calculated as outlined in Gedeon et al. ^70^. The code for carrying out the calculation is shared as <u>source/variableImportance.R</u> ^66^. The code uses the weights of the trained neural network to calculate the importance of each variable (input) used for classification. The output is scaled to 0 (least important) and 1 (most important). This was used to identify the importance of each gene used in zebrafish and human GERAS. The results were sorted in descending order for plotting. Additionally, the top 20 most important genes were obtained from the sorted list, and their relative importance calculated using the formula,

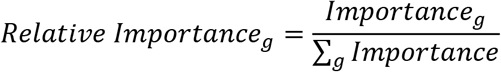

where *g* denotes an individual gene among the top 20. The disease association for each gene was obtained from DisGeNET database ^71^. From the database, an association with a score of greater than or equal to 0.2 was reported.

### Shiny implementation of GERAS predictor

To enable easy access to predictions using GERAS, a Shiny app was developed. The app is freely available at https://github.com/sumeetpalsingh/GERAS2017/shiny_GERAS_Tf.R^66^. The app provides a graphic-user interface (GUI) for users to make chronological age predictions using a pretrained GERAS model. The users can upload normalized counts, verify the uploaded data, and obtain predictions in a downloadable comma-separated (csv) file.

### Data availability

The raw datasets, along with tabulated count data and TPM normalized values, generated during the current study are available from GEO under accession number <u>GSE109881</u>, with the token number ixkzakssxnsjtaf. The data will be made public upon publication. Normalized read-counts for all human pancreatic samples used in the study are available at: https://sharing.crt-dresden.de/index.php/s/zcQ14AMGJAevokU, and codes for developing and testing GERAS are available at https://github.com/sumeetpalsingh/GERAS2017 ^66^. Please refer to README.md to navigate the Github folder. The authors welcome any requests for information on the raw data, data processing, GERAS development and utilization.

## Acknowledgements

We thank members of the Ninov lab for comments on the manuscript, members of Center for Regenerative Therapies Dresden (CRTD) fish, microscopy, sequencing and FACS facility for technical assistance. We are grateful to Priyanka Oberoi for illustrations.

## Authors’ Contributions

S.P.S. and Ankit Sharma (Google, N.Y.) conceptualized the project. S.P.S. and S.J. performed the zebrafish experiments. S.R., S.D. and A.E. performed single-cell sequencing. S.P.S., H.B., S.K., and G.Z. developed GERAS and its Shiny app. S.C., J.E.R. and N.N. provided critical feedback and advice. S.P.S., S.C. and N.N. wrote the manuscript. N.N. obtained key funding for the project. All authors read and approved the final manuscript.

## ADDITIONAL FILES

### Supplementary Figures (.pdf)

**Table S1: Variable Importance for zebrafish beta-cell GERAS (.xls).**

A table listing the 1000-input genes utilized by zebrafish beta-cell GERAS and their importance towards successful classification.

**Table S2: Genes negatively correlated with classification probability (.xls).**

For beta-cells from three-times-a-day fed animals, correlation analysis was performed. In the analysis, correlation coefficient was calculated between the probability to be classified in ‘Adolescent’ stage and gene expression. The genes were ranked in descending order of correlation coefficient. The table contains the genes in the bottom 5^th^ percentile.

**Table S3: Genes positively correlated with classification probability (.xls).**

For beta-cells from three-times-a-day fed animals, correlation analysis was performed. In the analysis, correlation coefficient was calculated between the probability to be classified in ‘Adolescent’ stage and gene expression. The genes were ranked in descending order of correlation coefficient. The table contains the genes in the top 5^th^ percentile.

**Table S4: Variable Importance for Human pancreatic GERAS (.xls).**

A table listing the 1000-input genes utilized by human pancreatic GERAS and their importance towards successful classification.

## References

1. Kowalczyk M. S. et al. Single-cell RNA-seq reveals changes in cell cycle and differentiation programs upon aging of hematopoietic stem cells. Genome Res. 25, 1860–1872 (2015).

2. Peters M. J. et al. The transcriptional landscape of age in human peripheral blood. Nat. Commun. 6, 8570 (2015).

3. Szilard L. On the nature of the aging process. Proc. Natl. Acad. Sci. U. S. A. 45, 30–45 (1959).

4. Vijg J. Somatic mutations and aging: a re-evaluation. Mutat. Res. Mol. Mech. Mutagen. 447, 117–135 (2000).

5. Pal S. & Tyler J. K. Epigenetics and aging. Sci. Adv. 2, e1600584–e1600584 (2016).

6. Barzilai, N., Huffman, D. M., Muzumdar, R. H. & Bartke, A. The Critical Role of Metabolic Pathways in Aging. Diabetes 61, 1315–1322 (2012).

7. López-Otín, C., Blasco, M. A., Partridge, L., Serrano, M. & Kroemer, G. The Hallmarks of Aging. Cell 153, 1194–1217 (2013).

8. Kenyon C. J. The genetics of ageing. Nature 464, 504–512 (2010).

9. Piper M. D. W. & Bartke A. Diet and Aging. Cell Metab. 8, 99–104 (2008).

10. Most, J., Tosti, V., Redman, L. M. & Fontana, L. Calorie restriction in humans: An update. Ageing Res. Rev. 39, 36–45 (2017).

11. Kõks S. et al. Mouse models of ageing and their relevance to disease. Mech. Ageing Dev. 160, 41–53 (2016).

12. Wang Y. & Navin N. E. Advances and Applications of Single-Cell Sequencing Technologies. Mol. Cell 58, 598–609 (2015).

13. Marco E. et al. Bifurcation analysis of single-cell gene expression data reveals epigenetic landscape. Proc. Natl. Acad. Sci. U. S. A. 111, E5643–50 (2014).

14. Haghverdi, L., Büttner, M., Wolf, F. A., Buettner, F. & Theis, F. J. Diffusion pseudotime robustly reconstructs lineage branching. Nat. Methods 13, 845–8 (2016).

15. Setty M. et al. Wishbone identifies bifurcating developmental trajectories from singlecell data. Nat. Biotechnol. 34, 637–45 (2016).

16. Trapnell C. et al. The dynamics and regulators of cell fate decisions are revealed by pseudotemporal ordering of single cells. Nat. Biotechnol. 32, 381–386 (2014).

17. Schiebinger G. et al. Reconstruction of developmental landscapes by optimal-transport analysis of single-cell gene expression sheds light on cellular reprogramming. bioRxiv doi:10.1101/191056 (2017). doi:10.1101/191056

18. Reid J. E. & Wernisch L. Pseudotime estimation: deconfounding single cell time series. Bioinformatics 32, 2973–2980 (2016).

19. Semrau S. et al. Dynamics of lineage commitment revealed by single-cell transcriptomics of differentiating embryonic stem cells. Nat. Commun. 8, 1096 (2017).

20. Chu L.-F. et al. Single-cell RNA-seq reveals novel regulators of human embryonic stem cell differentiation to definitive endoderm. Genome Biol. 17, 173 (2016).

21. Moignard V. et al. Decoding the regulatory network of early blood development from single-cell gene expression measurements. Nat. Biotechnol. 33, 269–276 (2015).

22. Ocone, A., Haghverdi, L., Mueller, N. S. & Theis, F. J. Reconstructing gene regulatory dynamics from high-dimensional single-cell snapshot data. Bioinformatics 31, i89–96 (2015).

23. Thattai M. & van Oudenaarden, A. Intrinsic noise in gene regulatory networks. Proc. Natl. Acad. Sci. 98, 8614–8619 (2001).

24. Raj, A., Peskin, C. S., Tranchina, D., Vargas, D. Y. & Tyagi, S. Stochastic mRNA synthesis in mammalian cells. PLoS Biol. 4, 1707–1719 (2006).

25. Battich, N., Stoeger, T. & Pelkmans, L. Control of Transcript Variability in Single Mammalian Cells. Cell 163, 1596–1610 (2015).

26. Stoeger, T., Battich, N. & Pelkmans, L. Passive Noise Filtering by Cellular Compartmentalization. Cell 164, 1151–1161 (2016).

27. Enge M. et al. Single-Cell Analysis of Human Pancreas Reveals Transcriptional Signatures of Aging and Somatic Mutation Patterns. Cell 171, 321–330.e14 (2017).

28. Martinez-Jimenez C. P. et al. Aging increases cell-to-cell transcriptional variability upon immune stimulation. Science. 355, 1433–1436 (2017).

29. Eldar A. & Elowitz M. B. Functional roles for noise in genetic circuits. Nature 467, 167–173 (2010).

30. Dueck, H., Eberwine, J. & Kim, J. Variation is function: Are single cell differences functionally important?: Testing the hypothesis that single cell variation is required for aggregate function. Bioessays 38, 172–80 (2016).

31. Kolodziejczyk, A. A., Kim, J. K., Svensson, V., Marioni, J. C. & Teichmann, S. A. The Technology and Biology of Single-Cell RNA Sequencing. Mol. Cell 58, 610–620 (2015).

32. Grün D. et al. Single-cell messenger RNA sequencing reveals rare intestinal cell types. Nature 525, 251–5 (2015).

33. Papalexi E. & Satija R. Single-cell RNA sequencing to explore immune cell heterogeneity. Nat. Rev. Immunol. (2017). doi:10.1038/nri.2017.76

34. Bader E. et al. Identification of proliferative and mature β-cells in the islets of Langerhans. Nature 535, 430–4 (2016).

35. Dorrell C. et al. Human islets contain four distinct subtypes of β cells. Nat. Commun. 7, 11756 (2016).

36. Singh S. P. et al. Different developmental histories of beta-cells generate functional and proliferative heterogeneity during islet growth. Nat. Commun. 8, 664 (2017).

37. Halpern K. B. et al. Single-cell spatial reconstruction reveals global division of labour in the mammalian liver. Nature 542, 352–356 (2017).

38. de Magalhães, J. P., Curado, J. & Church, G. M. Meta-analysis of age-related gene expression profiles identifies common signatures of aging. Bioinformatics 25, 875–881 (2009).

39. Glass D. et al. Gene expression changes with age in skin, adipose tissue, blood and brain. Genome Biol. 14, R75 (2013).

40. Gregg B. E. et al. Formation of a human α-cell population within pancreatic islets is set early in life. J. Clin. Endocrinol. Metab. 97, 3197–206 (2012).

41. Gunasekaran U. & Gannon M. Type 2 diabetes and the aging pancreatic beta cell. Aging (Albany. NY). 3, 565–75 (2011).

42. Ziegenhain C. et al. Comparative Analysis of Single-Cell RNA Sequencing Methods. Mol. Cell 65, 631–643.e4 (2017).

43. Oka T. et al. Diet-induced obesity in zebrafish shares common pathophysiological pathways with mammalian obesity. BMC Physiol. 10, 21 (2010).

44. Huang D. et al. The DAVID Gene Functional Classification Tool: a novel biological module-centric algorithm to functionally analyze large gene lists. Genome Biol. 8, R183 (2007).

45. Li, W., Hoffman, P. N., Stirling, W., Price, D. L. & Lee, M. K. Axonal transport of human a-synuclein slows with aging but is not affected by familial Parkinson’s disease-linked mutations. J. Neurochem. 88, 401–410 (2003).

46. Milde, S., Adalbert, R., Elaman, M. H. & Coleman, M. P. Axonal transport declines with age in two distinct phases separated by a period of relative stability. Neurobiol. Aging 36, 971–981 (2015).

47. Meynial-Denis D. Glutamine metabolism in advanced age. Nutr. Rev. 74, 225–36 (2016).

48. McIsaac, R. S., Lewis, K. N., Gibney, P. A. & Buffenstein, R. From yeast to human: exploring the comparative biology of methionine restriction in extending eukaryotic life span. Ann. N. Y. Acad. Sci. 1363, 155–70 (2016).

49. Hassa, P. O., Haenni, S. S., Elser, M. & Hottiger, M. O. Nuclear ADP-Ribosylation Reactions in Mammalian Cells: Where Are We Today and Where Are We Going? Microbiol. Mol. Biol. Rev. 70, 789–829 (2006).

50. Sikora, E., Kamińska, B., Radziszewska, E. & Kaczmarek, L. Loss of transcription factor AP-1 DNA binding activity during lymphocyte aging in vivo. FEBS Lett. 312, 179–82 (1992).

51. Ninov N. et al. Metabolic regulation of cellular plasticity in the pancreas. Curr. Biol. 23, 1242–1250 (2013).

52. Sakaue-Sawano A. et al. Visualizing spatiotemporal dynamics of multicellular cell-cycle progression. Cell 132, 487–98 (2008).

53. Segerstolpe, Å. et al. Single-Cell Transcriptome Profiling of Human Pancreatic Islets in Health and Type 2 Diabetes. Cell Metab. 593–607 (2016). doi:10.1016/j.cmet.2016.08.020

54. Kulas, J. A., Puig, K. L. & Combs, C. K. Amyloid precursor protein in pancreatic islets. J. Endocrinol. 235, 49–67 (2017).

55. van den Brink, S. C. et al. Single-cell sequencing reveals dissociation-induced gene expression in tissue subpopulations. Nat. Methods 14, 935–936 (2017).

56. Zeng C. et al. Pseudotemporal Ordering of Single Cells Reveals Metabolic Control of Postnatal α Cell Proliferation. Cell Metab. 25, 1160–1175.e11 (2017).

57. Aguayo-Mazzucato C. et al. β Cell Aging Markers Have Heterogeneous Distribution and Are Induced by Insulin Resistance. Cell Metab. 25, 898–910.e5 (2017).

58. Buenrostro J. D. et al. Single-cell chromatin accessibility reveals principles of regulatory variation. Nature 523, 486–490 (2015).

59. Lowsky, D. J., Olshansky, S. J., Bhattacharya, J. & Goldman, D. P. Heterogeneity in healthy aging. J. Gerontol. A. Biol. Sci. Med. Sci. 69, 640–9 (2014).

60. Jylhävä, J., Pedersen, N. L. & Hägg, S. Biological Age Predictors. EBioMedicine 21, 29–36 (2017).

61. Petkovich D. A. et al. Using DNA Methylation Profiling to Evaluate Biological Age and Longevity Interventions. Cell Metab. 25, 954–960.e6 (2017).

62. Belsky D. W. et al. Telomere, epigenetic clock, and biomarker-composite quantifications of biological aging: Do they measure the same thing? bioRxiv doi:10.1101/071373 (2016). doi:10.1101/071373

63. Stefan, N., Häring, H.-U. & Schulze, M. B. Metabolically healthy obesity: the low-hanging fruit in obesity treatment? lancet. Diabetes Endocrinol. (2017). doi:10.1016/S2213-8587(17)30292-9

64. Roberson L. L. et al. Beyond BMI: The ‘Metabolically healthy obese’ phenotype & its association with clinical/subclinical cardiovascular disease and all-cause mortality – a systematic review. BMC Public Health 14, 14 (2014).

65. Butler A. & Satija R. Integrated analysis of single cell transcriptomic data across conditions, technologies, and species. bioRxiv doi: 10.1101/164889 (2017). doi:10.1101/164889

66. Singh S. P. GERAS (GEnetic Referene for Age of Single-cell). (2017) https://github.com/sumeetpalsingh/GERAS2017.

67. Kim, D., Langmead, B. & Salzberg, S. L. HISAT: a fast spliced aligner with low memory requirements. Nat. Methods 12, 357–360 (2015).

68. Anders, S., Pyl, P. T. & Huber, W. HTSeq–a Python framework to work with high-throughput sequencing data. Bioinformatics 31, 166–169 (2015).

69. Castellazzi M. et al. Overexpression of c-jun, junB, or junD affects cell growth differently. Proc. Natl. Acad. Sci. U. S. A. 88, 8890–4 (1991).

70. Gedeon T. D. Data mining of inputs: analysing magnitude and functional measures. Int. J. Neural Syst. 8, 209–18 (1997).

71. Piñero J. et al. DisGeNET: a comprehensive platform integrating information on human disease-associated genes and variants. Nucleic Acids Res. 45, D833–D839 (2017).

